# Urban ecological connectivity as a planning tool for different animal species

**DOI:** 10.1101/2022.11.06.515356

**Authors:** Holly Kirk, Kylie Soanes, Marco Amati, Sarah Bekessy, Lee Harrison, Kirsten Parris, Cristina Ramalho, Rodney van der Ree, Caragh Threlfall

## Abstract

The application of ecological theory to urban planning is becoming more important as land managers focus on increasing urban biodiversity as a way to improve human welfare. City authorities must decide not only what types of biodiversity-focused infrastructure should be prioritized, but also where new resources should be positioned and existing resources protected or enhanced. Careful spatial planning can contribute to the successful return and conservation of urban nature by maximizing the contribution of green infrastructure to landscape connectivity. By using ecological connectivity theory as a planning tool, governments can quantify the effect of different interventions on the ease with which wildlife can move across the landscape. Here we outline an approach to a) quantify ecological connectivity for different urban wildlife species and b) use this to test different urban planning scenarios using QGIS. We demonstrate four extensions to the work by Deslaurier et al. (2018) and Spanowicz & Jaeger (2019) which improve the application of this method as a planning tool for local government:

- A step-by-step method for calculating effective mesh size using the open-source software QGIS.
- Conversion of the effective mesh size value (*m*_eff_) to a “probability of connectedness” (*P*_c_, for easier interpretation by local government and comparisons between planning scenarios).
- Guidance for measuring species-specific connectivity, including how to decide what spatial information should be included and which types of species might be most responsive to connectivity planning.
- Advice for using the method to measure the outcome of different urban planning scenarios on ecological connectivity.

## SPECIFICATIONS TABLE

|  |  |
| --- | --- |
| **Subject Area** | Environmental Science |
| **More specific subject area** | Urban Ecology |
| **Method name** | 'CBI Indicator 2' or 'effective mesh size, ($m_{eff}$)' |
| **Name and reference of original method** | Deslauriers, M.R., Asgary, A., Nazarnia, N., Jaeger, JAG. 2018. Implementing the connectivity of natural areas in cities as an indicator in the City Biodiversity Index (CBI) (+ Corrigendum). *Ecological Indicators* **94** (2) 99-113<br>Spanowicz, A.G. & Jaeger, J.A.G. 2019. Measuring landscape connectivity: On the importance of within-patch connectivity. *Landscape Ecology* **34** (10) 2261-2278 |
| **Resource availability** | QGIS version 2.18 or later |

## Method details

### Rationale

To guarantee their survival and long-term population viability, species need to be able to move around a landscape and access resources (e.g. food, shelter or suitable mates) (Hanski, 1999; Taylor et al., 1993). Ecological connectivity refers to the ability of species to move through a landscape, that considers how different landscape attributes, such as habitat patches, facilitate or impede animal movement (Taylor et al., 1993). Quantifying ecological connectivity has become an important component in conservation planning, leading to the development of a several different methods for measuring the connectedness of a landscape (Brooker et al., 1999; Bunn et al., 2000; Moilanen & Nieminen, 2002; Tischendorf & Fahrig, 2000; Urban & Keitt, 2001). More recently, a focus on conserving urban diversity has led to cities considering ecological connectivity as part of their urban planning process (Braaker et al., 2017; Casalegno et al., 2017; LaPoint et al., 2015; Lookingbill et al., 2022). Urban environments are often highly fragmented landscapes, in which most habitat suitable for animal species, such as patches of remnant, semi-natural and managed vegetation, are surrounded by a matrix of residential, commercial and transportation land-uses that prioritise human activities. A high level of ecological connectivity within an urban landscape enables animals to move between patchy resources and allows post-breeding dispersal, maintaining gene flow and population viability (Braaker et al., 2014, 2017; Mimet et al., 2020; Ossola et al., 2019; Visscher et al., 2018). In contrast, a low level of ecological connectivity prevents the movement of individuals and genes, potentially leading to reduced genetic diversity, inbreeding depression and ultimately local extinction (Hanski, 1999; Keeley et al., 2017; LaPoint et al., 2015; Moilanen & Hanski, 2001; Teitelbaum et al., 2020).

Here we present an extension to the method for quantifying ecological connectivity initially proposed by Deslauriers et al. (2018) for the City Biodiversity Index (CBI, sometimes known as the Singapore Index), Indicator 2 (*IND2*_CBI_impr_). Deslaurier et al. (2018) use effective mesh size (*m*_eff_), a geometric measure for estimating landscape fragmentation, based on “the probability that two randomly chosen locations in the landscape are connected and not separated by barriers” (Jaeger, 2000; Spanowicz & Jaeger, 2019). Effective mesh size (*m*_eff_) is a measure of the area of habitat that can be accessed by an individual organism when dropped at random into the landscape (Spanowicz & Jaeger, 2019). This is a useful method for quantifying urban connectivity, as it is easily calculated with commonly available GIS software (such as ArcGIS) and a spreadsheet program (such as Microsoft Excel). It also generates a measure of area, rather than an index, which is more easily interpreted by the multiple, non-ecological disciplines involved in urban planning. The use of raw spatial data inputs rather than arbitrary values for model parameterization is another advantage of effective mesh size over other measures of ecological connectivity (Lookingbill et al., 2022). Effective mesh size allows comparison of connectivity between different places (e.g. municipalities or local government areas) even if they differ in terms of total area. Deslaurier et al. (2018) and Spanowicz & Jaeger (2019) both provide a step-by-step method for calculating connectivity using ArcGIS and demonstrate how this method can be applied to quantify and compare ecological connectivity in real (Montréal and Lisbon, Deslaurier et al. 2018), and theoretical landscapes (Spanowicz & Jaeger, 2019). Both studies used generalised values for the three main input parameters used to measure landscape connectedness (habitat area, barriers to movement and interpatch threshold distance).

The method described by Deslauriers et al. (2018) & Spanowicz & Jaeger (2019) uses very broad definitions of habitat, barrier and interpatch threshold distance. This means that, aside from being used as a standard benchmark comparison with other municipalities, this structural connectivity measure is harder to interpret in real terms from the perspective of animal species living in a city. By including a functional approach (see Casalegno et al., 2017; Ersoy et al., 2019; Grafius et al., 2017; Mimet et al., 2013; Nor et al., 2017; Vogt et al., 2009) and choosing more realistic definitions for the model parameters, the method we propose can be tailored to a single species or species group. This gives a better estimate of how connected the landscape is for specific animals, enabling local governments to plan for a single target species or group. Our method extension details: 1) a step-by-step method for calculating effective mesh size (*m*_eff_) using the open-source software program QGIS, including the conversion to a “probability of connectedness” (*P*_c_); 2) guidance for measuring species-specific connectivity, including how to decide what spatial information should be included in the connectivity model; and 3) examples of using the measure to compare urban planning scenarios.

### Calculating effective mesh size using QGIS

Effective mesh size (*m*_eff_) quantifies landscape connectivity as the “average area of habitat accessible to an individual dropped randomly anywhere in the landscape” (Deslauriers et al., 2018; Moser et al., 2007; Spanowicz & Jaeger, 2019). Connectivity is measured by quantifying the relationship between the total area of habitat available in the landscape and the degree to which this total area is fragmented, either by distance (between individual habitat patches) or by barriers to movement, such as a road network. The workflow for calculating effective mesh size first requires the identification of habitat patches. These patches are then determined to be either connected or not connected to one another. For a habitat patch to be classified as connected to another patch or patches it must be 1) within the threshold distance for inter-patch connectivity and 2) not separated by a barrier to movement. Groups of connected patches are identified using the method, their area determined and then used to calculate effective mesh size and probability of connectedness.

The method for calculating effective mesh size, *m*_eff_ (illustrated in Figure 1) has six main steps:

**Figure 1.**
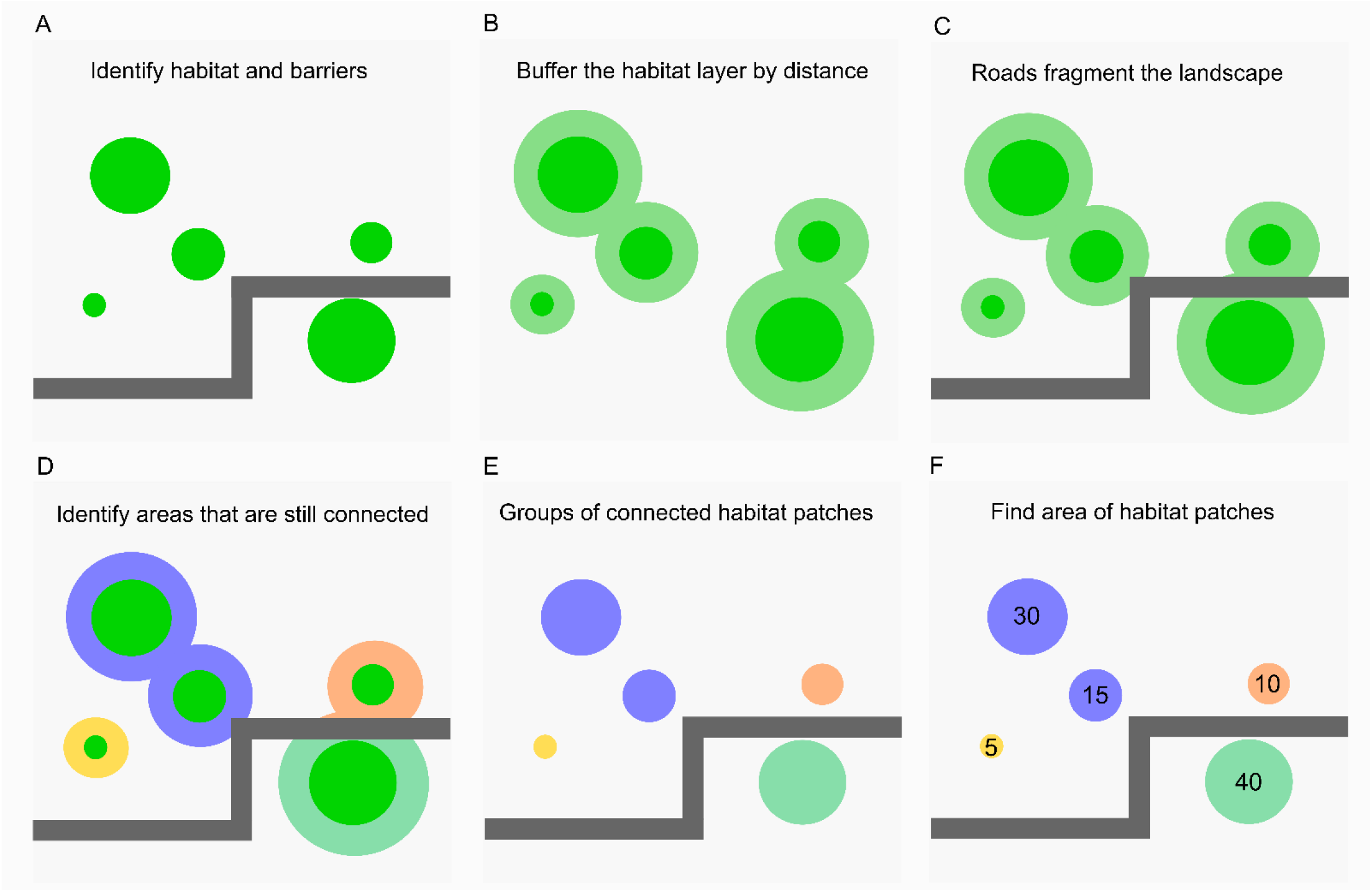
Schematic representation of how urban ecological connectivity is measured using effective mesh size (m_eff). In this example, the landscape contains four patches of habitat (A). Buffering these habitat patches by a fixed distance determines which patches are connected (B). However, the road network provides additional barriers to movement, increasing fragmentation (C). In this example, the two blue patches are part of the same connected area (D and E). The area of each individual vegetation patch is then calculated (F). These values are used to calculate the connectivity index using equation 1: sum of all patch areas = 45+10+5+40 = 100 (note that the two patches that make up the blue connected area are summed together); sum of all patch areas squared = 2025+100+25+1600 = 3750; m_eff = 3750/100 = 37.5; P_c= 37.5/100 = 0.375.

1. Identify land uses within the landscape that can be considered ‘habitat’ and which land uses are ‘barriers’ to movement (Figure1A).
2. Buffer around areas of ‘habitat’ to identify which patches are ‘connected’, i.e. are close enough together that the gap between them is not impassable for the species in question. E.g. 50m buffer around habitat patches will identify the patches that are more than 100m apart (Figure 1B).
3. Remove any ‘barriers’ from the buffered layer (this creates a ‘fragmentation geometry’). E.g. if roads wider than 10m are considered a barrier to movement then habitat patches either side of these roads are not ‘connected’, even if they are within 100m of each other (Figure 1C)
4. Identify the buffered areas that are still connected (‘connected areas’, Figure 1D).
5. Identify which habitat patches belong in each of the connected areas (‘connected patches’, Figure 1E).
6. Calculate the total area of each set of connected habitat patches (Figure 1F). These are fed into the effective mesh size equation (1).

Steps 1 to 5 describe how to determine which patches of habitat are “connected” according to how far apart the patches are and whether any barriers to movement exist between them. This is the spatial part of the analysis which requires GIS software. The results of this part of the analysis are then exported as purely numeric values and used in Step 6, which is where the relationship between total habitat area and connected habitat area is quantified using effective mesh size. The equation for calculating effective mesh size (*m*_eff_) is:

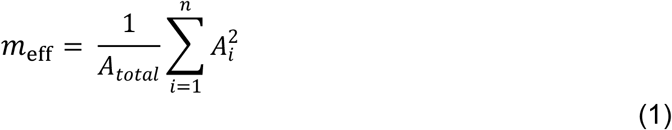

Where *n* = the number of groups of connected habitat patches, *A*_*total*_ = the total area of all habitat patches and 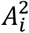 = squared area of each of the *n* groups (*i* = 1, …, *n*) of connected habitat patches. (Spanowicz & Jaeger, 2019). The formula can be simplified to the following:

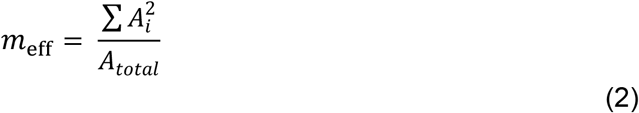

Effective mesh size is a measure of area (e.g., m^2^ or Ha) which increases as the habitat patches within the landscape become more connected (Deslauriers et al. 2018). This sometimes results in very large values that can make it harder to interpret, especially for non-specialists. With this in mind, we also suggest calculating a “probability of connectedness” (*P*_c_), or the “probability that two individuals dropped into the landscape will be in the same connected area of habitat”. Once effective mesh size has been found, the probability of connectedness (*P*_c_) can be calculated simply using the following method:

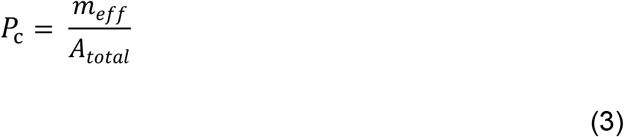

This is equivalent to Jaeger’s “degree of coherence” (*C*, Jaeger, 2000):

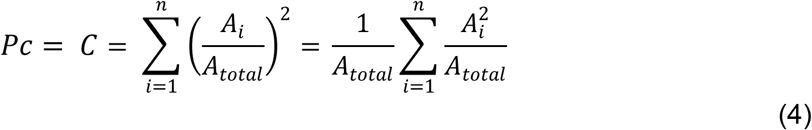

The probability of connectedness scales effective mesh size to a value between 0 and 1, where 1 would be a completely connected landscape for the individuals in question. Table 1 details the step-by-step methodology for calculating effective mesh size and probability of connectedness using the open-source GIS software QGIS (QGIS.org, 2021) and MS Excel spreadsheet software. See the ‘*Method Application*’ section below for an illustration of how effective mesh size and probability of connectedness were measured for species in the City of Melbourne, Australia.

**Table 1.** (Over page) Step-by-step instructions for calculating urban ecological connectivity in QGIS, from initial data layer preparation, through the spatial analysis to determine connected areas and final mathematical calculation of connectivity. *An additional step may be required here to remove any habitat patches that exist ON TOP of the barriers (e.g. roadside trees, median strips). Do this using “Geoprocessing > Difference” Input layer = habitat layer from step 1.1; Difference layer = barrier layer from step 2.1 or 2.2. ** When using this method to compare the connectivity before and after the addition or removal of habitat the original/existing total area of habitat (rather than the new area) should be used as the denominator in the effective mesh size calculation.

| Analysis step | Method in QGIS/Excel | Rationale |
| --- | --- | --- |
| **Prepare habitat layers** |  |  |
| 1.1 Unify habitat layers | Unify the habitat layers (you may also like to dissolve after each union to limit file size)<br>Geoprocessing tools > Union: Input the two layers you wish to combine | If using more than one habitat layer, combine them to form one layer. E.G. grass AND trees. Do this step as many times as needed to combine 3 or more layers. |
| 1.2 Buffer habitat layers | Buffer & dissolve habitat layers.<br>Geoprocessing > Fixed distance buffer: distance value is half the dispersal distance for your species. Select "dissolve result" | Habitat patches are classed as "connected" if less than the "dispersal distance" apart. Patches closer than this will be grouped together. |
| **Prepare barrier layers** |  |  |
| 2.1 Unify barrier layers | Unify the barrier layers (you may also like to dissolve after each union to limit file size)<br>Geoprocessing tools > Union: Input the two layers you wish to combine. | Combine the different types of barrier together to make single layer. E.G. roads AND buildings |
| 2.2 Buffer barrier layers (optional) | Buffer & dissolve barrier layers<br>Geoprocessing tools > Fixed distance buffer: distance value chosen to represent "edge effects". Select "dissolve result" | Barriers such as roads may have an "edge effect" on the habitat around them. Distance value used in City Biodiversity Index is 7.5m. |
| **Create fragmentation geometry** |  |  |
| 3.1 Erase barriers from the buffered habitat layer | Input layer = buffered habitat layer 1.2<br>Difference layer = prepared barrier layer 2.1 or 2.2<br>Geoprocessing tools > Difference. | Barriers can prevent movement between habitat patches, even if they are within the normal dispersal distance for a species. This provides an additional level of fragmentation. |
| **Identify connected areas \*** |  |  |
| 4.1 Find the remaining connected areas | Input layer = fragmentation geometry 3.1<br>Geometry tools > Multipart to singleparts. | Identifies each different connected area (group of habitat patches). |
| 4.2 Identify connected areas | Right click on the layer created in step 4.1, open attribute table and toggle editing on. Open "field calculator" and "create new field" using "row_number" function. | Gives each connected area that a unique number in a new field. Call the new field something sensible like "patchID" or "area_number" |
| **Identify connected habitat patches** |  |  |
| 5.1 Intersect original habitat layer with identified connected areas | Input layer = layer created in step 4.2<br>Intersect layer = habitat layer from step 1.1 (combined habitat types)<br>Geoprocessing > Intersection | Links each original habitat patch with its related connected area so that connected patches of habitat are grouped together with the same identifier (generated in step 4.2). |
| **Calculate area and export to .csv** |  |  |
| 6.1 Calculate area of each patch | Right click on the layer created in step 5.1. Open attribute table and toggle editing on. Open "field calculator" and "create new field" using "area" function. Use the maximum level of precision possible (e.g. 10 dp & 'decimal number' rather than 'integer') | This step calculates the actual area of each connected habitat patch. A patch is a collection of smaller pieces of habitat that are classed as connected due to their proximity/lack barriers |
| 6.2 Export attribute table to ".csv" | Right click the layer from step 6.1 and select "save as". Make sure to select .csv file type | The final calculation is done in Excel |
| **Connectivity calculation \*\*** |  |  |
| 7.1 Calculation of (area)<sup>2</sup> | Spreadsheet should include columns: patch ID and area. Calculate square of patch area for each habitat patch (i.e. for the whole column of data). | Open Excel or any spreadsheet tool. The first part of the calculation requires the generation of a new column, area<sup>2</sup>. |
| 7.2 Calculate effective mesh size | $m_{\text{eff}} = \text{SUM of squared patch areas} / \text{SUM of original habitat area}$ | This is the measure of connected area available |
| 7.3 Calculate probability of connectance | $P_c = m_{\text{eff}} / \text{SUM of original habitat area}$ | This is also known as the "degree of coherence" |

### Species-specific ecological connectivity

Many recent urban connectivity studies focus on assessing the overall connectedness of vegetation within a landscape using a set of generalised parameters to determine if two patches of vegetation are “connected” or not (e.g. Casalegno et al., 2017; Liu et al., 2022; Lookingbill et al., 2022; Ossola et al., 2019). For example, the City Biodiversity Index specifies 100 m as the threshold distance to determine if vegetation patches are connected and roads of >15 m in width are classified as barriers to movement (determining the fragmentation geometry, Chan et al. 2014). While these generalised parameters are useful for an overall assessment of connectivity, realistically these parameters are too broad to help with decision making for many species groups. For example, many bird species are able to disperse or commute across inter-patch distances of more than 100m (Shanahan et al., 2011; Silva et al., 2020; Tremblay & St. Clair, 2011), while amphibian species are likely to be impeded by traffic-heavy roads (Charry & Jones, 2009; Hamer, 2016; Jacobson et al., 2016). To measure connectivity more realistically for different species groups it is important to account for behaviour and ecology (Ersoy et al., 2019; Grafius et al., 2017; Lookingbill et al., 2022; Mimet et al., 2020; Teitelbaum et al., 2020; Tischendorf & Fahrig, 2000). Each species within a community has different habitat requirements and movement capabilities. This means the definition of “connectedness” changes depending on the species or group that is being considered. By considering the distinct resource requirements and movement capability of a focal animal species the functional landscape connectivity for that species can be measured (Baguette et al., 2014; Tischendorf & Fahrig, 2000; Ziółkowska et al., 2014).

### Choosing species for modelling

When using effective mesh size for urban ecological connectivity planning it is helpful to choose a focal species or group of species for modelling. A focal species approach not only permits clearer decision making when it comes to management actions (Mata et al., 2020), but also allows better definition of the habitat being measured and its level of fragmentation within a city, which will vary depending on the species being considered (Ziółkowska et al., 2014). Selecting focal species will not only account for variability in how connectedness is defined for different groups but allows the resulting connectivity measures to guide more specific management actions, such as increasing flowering shrub cover in parks or creating artificial nesting hollows. Choosing focal species for connectivity modelling may depend on the needs of the land manager, some may already have species that are being considered as conservation priorities or targets for restoration. Where focal species have not been pre-determined, several factors should be considered when choosing focal species for connectivity modelling:

- Species distributions and local populations – species should already be present within the local area, or even just outside. Species with distributions a long distance from the area being considered are unlikely to benefit from local actions, particularly if beyond their usual movement patterns.
- Response of the focal species – focal species for ecological connectivity planning in urban spaces should be species that are able to persist in urban environments under the right conditions, meaning that they are likely to respond well to management actions that aim to improve their ability to move across the landscape
- Planned actions – land managers may wish to test the effect that different management actions will have on urban ecological connectivity. Choosing species that will be impacted (positively or negatively) by those actions will be useful. For example, increasing tree canopy may not have a big effect on connectivity for terrestrial reptiles

Stakeholder consultation processes can be a highly effective way to choose focal species. Involving a diverse range of local stakeholders in the decision process may also increase the support for connectivity-focused management actions and even create more positive human-nature interactions (Apfelbeck et al., 2019). Expert elicitation, such as following the IDEA protocol could also be a useful approach (Hemming et al., 2018). Focal species decision workshops can then be coupled with expert input on the appropriate habitat resources, movement ability and potential movement barriers used to determine connectedness (see below). Alternatively, species selection frameworks for general urban conservation decision-making can be used to choose appropriate species for connectivity modelling (Apfelbeck, 2020; Garrard et al., 2018; Mata et al., 2020; Parris et al., 2018).

Predicting ecological connectivity is data intensive, as it requires detailed information about the target species’ movement abilities, habitat preferences and potential barriers to dispersal. This information is often not known or unavailable, especially for urban environments. To represent the ecological connectivity of a broad range of animal species within a municipality a species group approach may be useful. By considering groups or guilds of animals that share similar dispersal abilities and habitat requirements the results of connectivity modelling can be extrapolated across a few different species, e.g. all native bees or all amphibians.

### Selecting appropriate habitat, barrier and distance definitions

Measuring connectivity using effective mesh size and probability of connectedness requires each part of the landscape to be characterised based on its potential to provide habitat and impede or promote movement for the targeted species or species groups. Information on how a focal species moves through the landscape is therefore important. Tracking or mark-recapture data are particularly useful to inform and help generalise how a species uses different landscape features and elements, because these data can then be used to map connectivity at broader landscape or regional scales (Calabrese & Fagan, 2004; Tischendorf & Fahrig, 2000). However, collecting information on species movement behaviour requires extensive field research, and consequently, empirical data is often scarce (Pe’er et al. 2011; La Point et al. 2015; Lookingbill et al., 2022). The most important estimate for calculating effective mesh size and probability of connectedness is the distance which an individual can move across a gap in habitat, or its ‘gap-crossing ability’. Depending on the type of species being considered, this distance threshold might be determined from the species’ post-breeding dispersal distance, or the area an individual covers during daily foraging movements, sometimes called a home-range estimate. In general, post-breeding dispersal distances tend to be larger than daily-home range movements, but both are important processes for species persistence. In the absence of empirical movement data specific to the species and landscape of focus, estimates for how the focal species might move across the landscape can be derived either from similar species or groups within the literature, or by consulting with species experts, perhaps using an expert elicitation framework. Where a broad range of movement distances have been recorded for a species or a species group (such as daily foraging movements and post-breeding dispersal) we recommend measuring connectivity at least twice using the minimum and maximum distances. This will help to account for uncertainty and produce a better understanding of opportunities for improving/protecting ecological connectivity for that species.

Selection of appropriate spatial data layers is also important and will vary from one municipality to another depending on availability. Data layers that identify the location of existing habitat should include vegetation cover, such as tree canopy extents, understorey vegetation, location of managed garden beds, managed turf and areas classified as parks or reserves. If an aquatic species is chosen for modelling then including permanent waterways and waterbodies is necessary, and depending on the species considered, ephemeral wetland resources such as rain gardens, swales and drainage lines can also be classed as potential habitat. Identifying habitat patches requires careful consideration of the resources used by the species and how these may be represented across several sources of spatial data. For example, to create a habitat layer for a nectivorous bird that uses riparian zones, tree canopy and flowering shrubs will require a combination of several data layers. Table 1 details the QGIS steps for combining multiple habitat layers together in preparation for the effective mesh size calculation. It is important to note that spatial vector data for habitat areas can often be very complex, especially if derived from LiDAR. The running time of the connectivity analysis can be decreased by reducing the complexity of the habitat layer by unifying or adjacent boundaries in the polygon layer (in QGIS you can used the “dissolve” function) or simplifying the layer during data preparation. Data layer choice should be supported by the existing ecological literature or consultation by species experts where needed. Examples of how several data layers were combined to identify existing habitat patches for seven species groups in the City of Melbourne, Australia, can be found in Table 2.

**Table 2.**
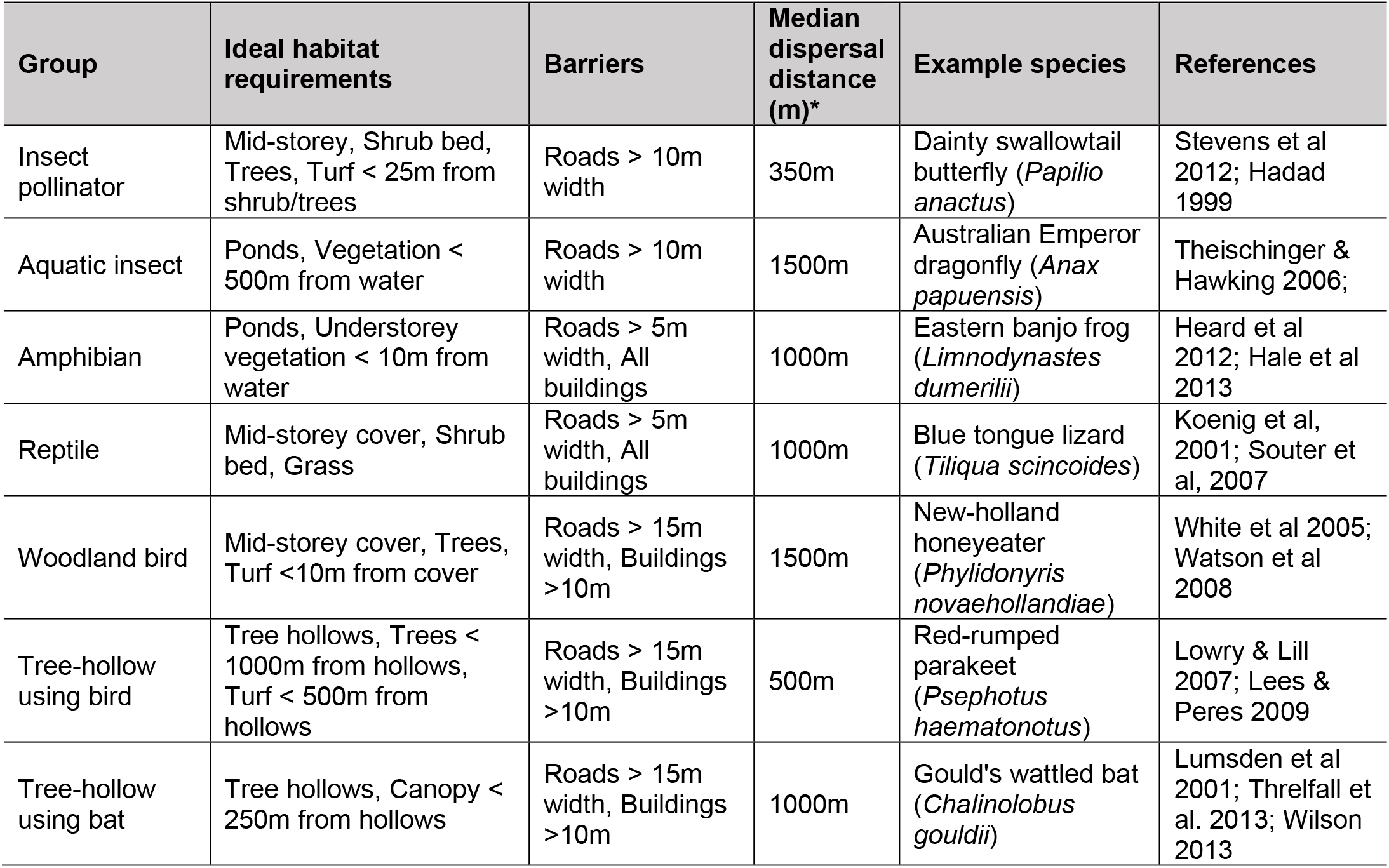
Habitat, barrier and movement threshold definitions used to estimate ecological connectivity across the City of Melbourne, Australia for seven animal groups. *Gap crossing ability or the smallest mean home range/minimum mean movement distance.

| Group | Ideal habitat requirements | Barriers | Median dispersal distance (m)* | Example species | References |
| --- | --- | --- | --- | --- | --- |
| Insect pollinator | Mid-storey, Shrub bed, Trees, Turf < 25m from shrub/trees | Roads > 10m width | 350m | Dainty swallowtail butterfly (*Papilio anactus*) | Stevens et al 2012; Hadad 1999 |
| Aquatic insect | Ponds, Vegetation < 500m from water | Roads > 10m width | 1500m | Australian Emperor dragonfly (*Anax papuensis*) | Theischinger & Hawking 2006; |
| Amphibian | Ponds, Understorey vegetation < 10m from water | Roads > 5m width, All buildings | 1000m | Eastern banjo frog (*Limnodynastes dumerilii*) | Heard et al 2012; Hale et al 2013 |
| Reptile | Mid-storey cover, Shrub bed, Grass | Roads > 5m width, All buildings | 1000m | Blue tongue lizard (*Tiliqua scincoides*) | Koenig et al, 2001; Souter et al, 2007 |
| Woodland bird | Mid-storey cover, Trees, Turf <10m from cover | Roads > 15m width, Buildings >10m | 1500m | New-holland honeyeater (*Phylidonyris novaehollandiae*) | White et al 2005; Watson et al 2008 |
| Tree-hollow using bird | Tree hollows, Trees < 1000m from hollows, Turf < 500m from hollows | Roads > 15m width, Buildings >10m | 500m | Red-rumped parakeet (*Psephotus haematonotus*) | Lowry & Lill 2007; Lees & Peres 2009 |
| Tree-hollow using bat | Tree hollows, Canopy < 250m from hollows | Roads > 15m width, Buildings >10m | 1000m | Gould's wattled bat (*Chalinolobus gouldii*) | Lumsden et al 2001; Threlfall et al. 2013; Wilson 2013 |

Barriers to movement are a critical consideration in connectivity modelling. Potential barriers in urban landscapes include transport corridors, tall buildings (such as skyscrapers) or wide rivers. Road networks are the most common movement barrier type. However, the extent to which a road restricts animal movement depends on the road width, traffic volume and species mobility. Furthermore, in some cases roadside vegetation allows road networks to function as corridors of habitat, rather than simply barriers. Deciding which land use classes constitute a barrier to movement for each focal species has perhaps the most impact on the outcome of the connectivity model. This is because the effective mesh size measure of ecological connectivity uses a binary definition to determine whether a barrier exists between two patches (visualised in Figure 1C and 1D and for an insect pollinator species in Figure 2C). Therefore, capturing the complexity of whether a land use is a complete barrier to movement, or more of a deterrent, can be difficult, especially in the absence of urban-specific animal behaviour research to underpin these decisions. This is perhaps the biggest limitation of the effective mesh size approach for measuring connectivity when compared to more complex methods such as least cost path or circuit theory (Balbi et al., 2019; Dickson et al., 2019; Grafius et al., 2017). These other methods take a more subtle approach, allowing a gradient of “movement” across different land use types. However, this approach requires additional model parameterisation which is often left up to professional judgment in the absence of empirical data on species movement and behaviour within urban areas (Lookingbill et al., 2022). Our species- or taxon-specific approach provides an intermediate solution: while the final model still uses an binary definition of connectivity across movement barriers (if a barrier exist between two habitat patches then they are not connected), the decision of which a land uses are classed barriers is determined by what is known about the species’ ecology. Table 2 shows how potential barrier land use layers were defined for seven species groups in the City of Melbourne.

**Figure 2.**
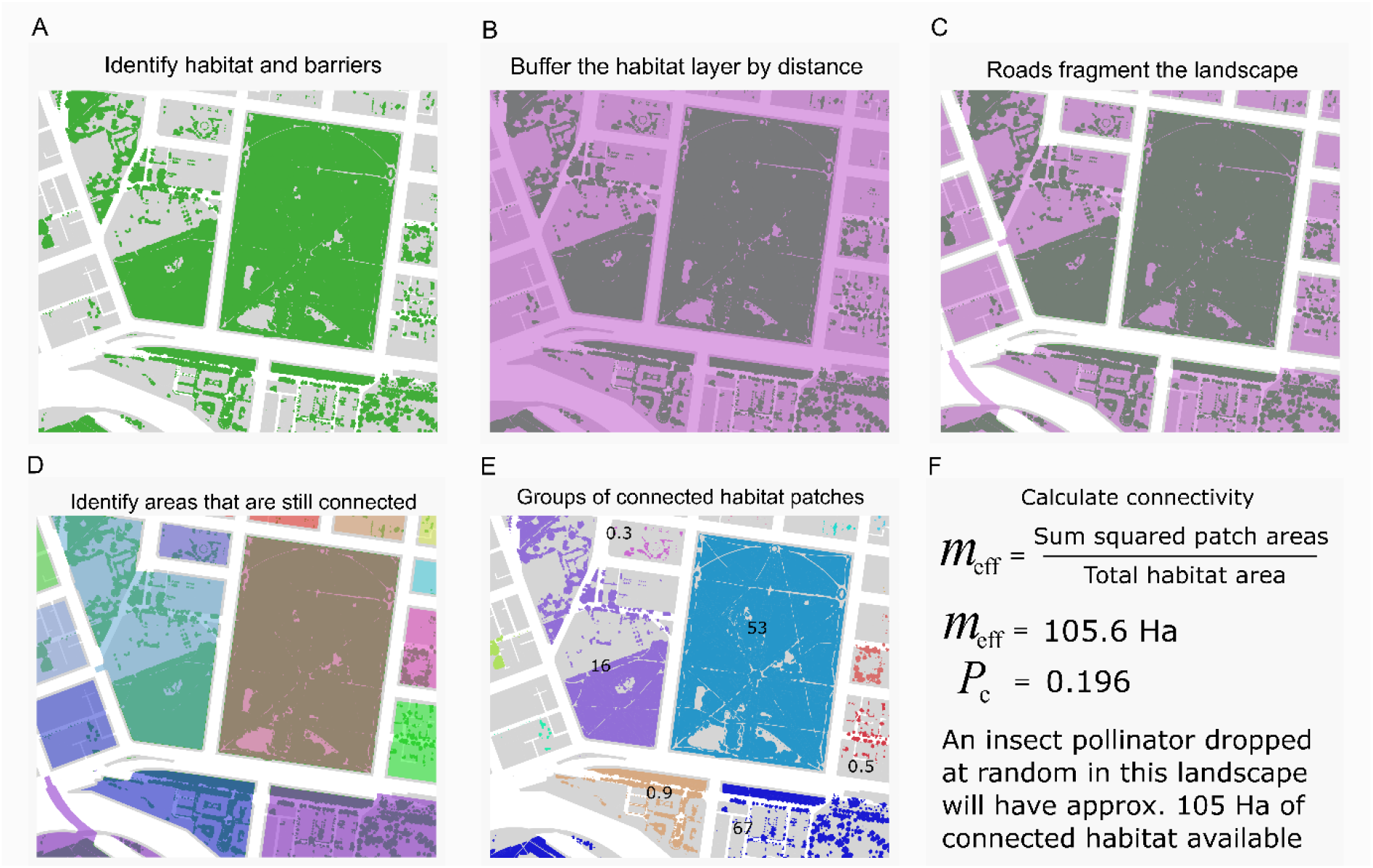
Example of the how existing ecological connectivity for an insect pollinator is measured using effective mesh size across a small area within the City of Melbourne, Australia. In this example, the landscape contains multiple small patches of habitat and a few larger urban parks containing understorey vegetation, coloured green (A). Buffering these habitat patches by 350m determines which patches can be reached by the flying insect (B). The roads wider than 10m provide additional barriers to movement, increasing the level of fragmentation (C). The connected areas are then identified, with different colours used to show this (D), which allows the groups of connected habitat patches to be identified (and coloured accordingly for visualisation purposes, E).

### Method validation - example application in real-life planning scenarios

Here we demonstrate how our method can be used to 1) determine the existing ecological connectivity of a highly urbanized landscape, and 2) explore the impact of two planning scenarios on the ecological connectivity of the same landscape. This work formed part of a connectivity plan developed with the City of Melbourne council. The City of Melbourne municipality is an inner-city area of approximately 37.7 km^2^ within the coastal city of Greater Melbourne, Victoria, Australia (figures 3 and 4 show the extent of the local government boundary).

**Figure 3.**
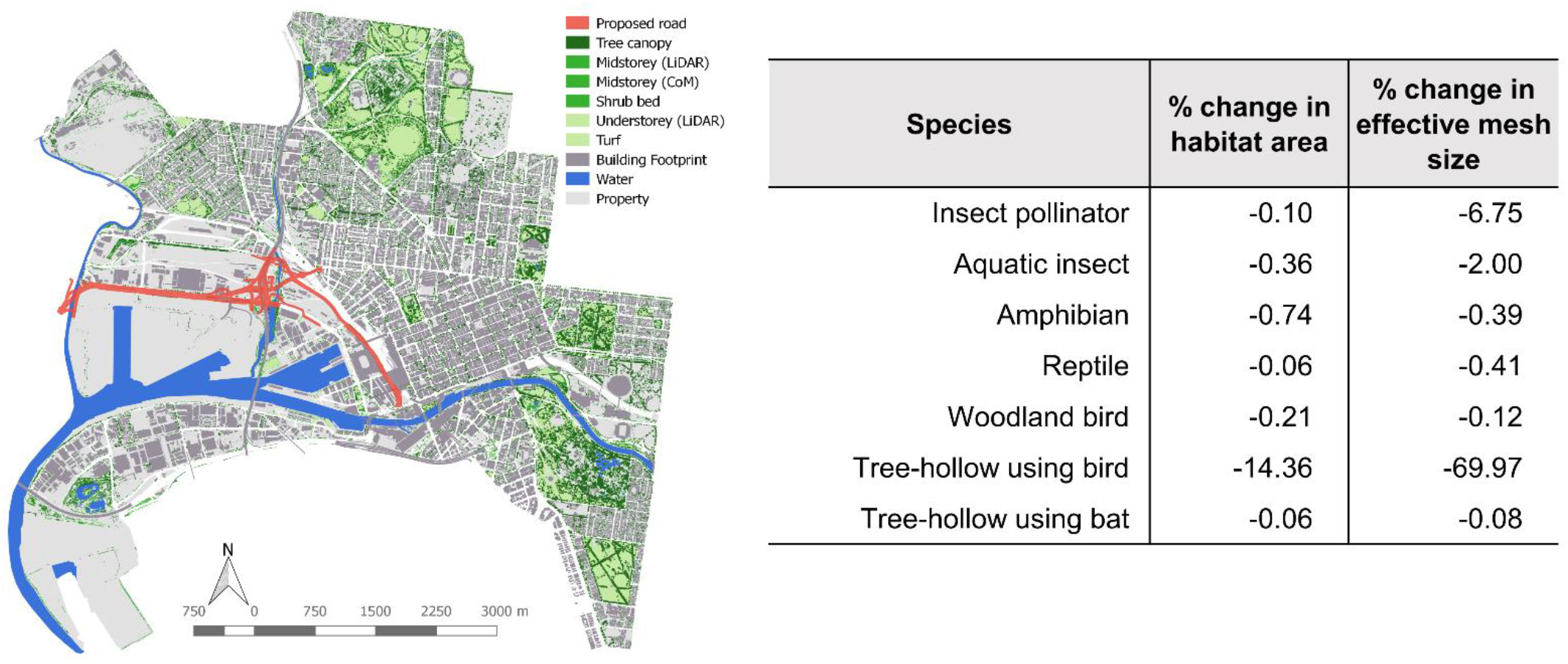
Visualisation showing an example of a planning scenario with a potential negative impact on ecological connectivity. The map shows an overview of the City of Melbourne with existing habitat patches (green) and a hypothetical new road development red. Habitat occurring underneath the new road network was removed and the effective mesh size before and after this simulation measured. The table shows the decrease in effective mesh size for most species, accompanied by a reduction in total area of habitat available.

**Figure 4.**
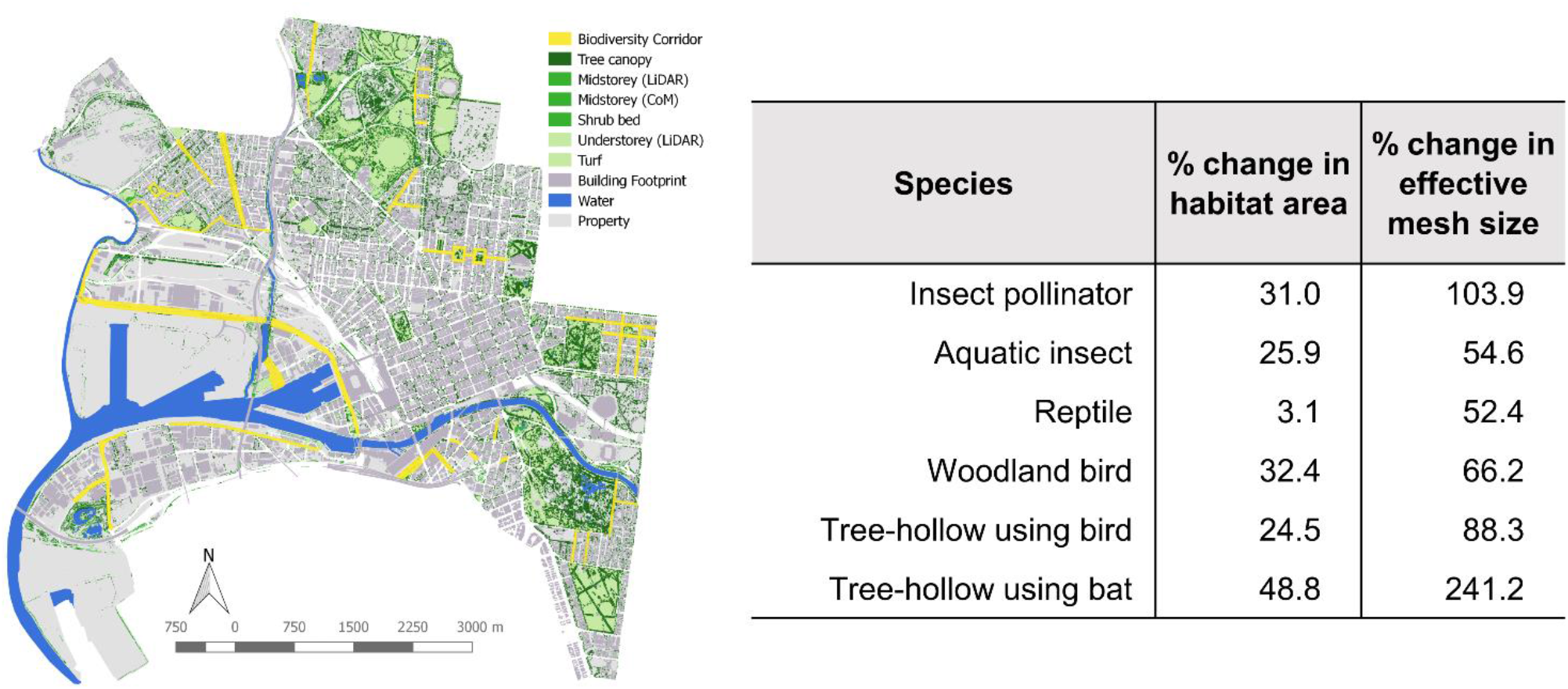
Visualisation showing an example of a planning scenario with a potential positive impact on ecological connectivity. The map shows the potential results of an urban forest planting strategy that aims to improve biodiversity within the City of Melbourne. Biodiversity corridors (yellow) should have full tree canopy cover in the future. Existing vegetation is shown in green. The contribution of these additional areas of habitat to the landscape were measured by calculating effective mesh size before and after. The table shows an increase in effective mesh size for all species modelled, accompanied by an increase in the total area of habitat available and a reduction in the number of connected areas as more patches of habitat have become connected. Note that the ‘Amphibian’ species group was not included in this scenario as the modelled habitat change did not include resources for this species.

Existing ecological connectivity in the City of Melbourne was quantified for seven animal groups:

- Insect pollinators – species where the adult stages depend on flowering vegetation as a food resource (e.g. blue-banded bee, *Amegilla chlorocyanea*);
- Aquatic insects – species dependent on waterbodies for larval life stages but that are also able to move overland as flying adults (e.g. blue skimmer dragonfly, *Orthetrum caledonicum*);
- Amphibians – species that depend on waterbodies for reproduction and are limited in overland dispersal (e.g. spotted marsh frog, *Limnodynastes tasmaniensis*);
- Reptiles – species that depend on adequate ground cover for refuge, including leaf litter, rocks and coarse woody debris (e.g. eastern blue-tongued lizard, *Tiliqua scincoides*);
- Woodland birds – species that depend on dense or complex mid-storey vegetation for nesting and resources (e.g. superb fairy-wren, *Malurus cyaneus*);
- Tree-hollow using birds – species that depend on tree hollows for reproduction and fly above and below the tree canopy during the day (e.g. red-rumped parrot, *Psephotus haematonotus*); and
- Tree-hollow using bats – species that depend on tree hollows for refuge and reproduction and move within or closely associated to the tree canopy during the night (e.g. Gould’s wattled bat, *Chalinolobus gouldii*).

These groups cover the habitat range and movement abilities of many species within the City of Melbourne, including multiple taxa (e.g. insects, amphibians, reptiles, birds and mammals), and multiple levels of habitat structure (e.g. water, ground cover, mid-storey and tree canopy). Each group also contains species that are charismatic and likely to garner public support for conservation actions. We used the primary literature and consultation with a panel of experts (workshop held on 11th August 2017; see acknowledgements) to identify movement ability or median dispersal distance, potential barriers and habitat requirements within the landscape for each group. Further details on the characteristics of each group are provided in Table 2. To model the landscape, GIS data were gathered in the form of ESRI shapefiles. This included LiDAR data collected in 2014, which covered both public land and private land within the municipality that is not managed by the City of Melbourne council. LiDAR data showed the extent of three vegetation classes grouped by height: vegetation under 50 cm, vegetation between 50 cm and 300 cm and vegetation greater than 300cm. These layers were supplemented with spatial data from the City of Melbourne, providing extra information about public land management, including parks, reserves, waterbodies, road medians, street trees and garden beds. GIS data layers describing the road, rail and building footprints were classed as barrier land uses for most species and determined the level of fragmentation within the landscape.

Figure 2 illustrates the spatial analysis steps for calculating the effective mesh size of existing insect pollinator habitat within the City of Melbourne. The first panel visualizes habitat in and around an urban park (Fitzroy Gardens, East Melbourne) and the surrounding road network. The later panels in Figure 2 illustrate the spatial output that can be generated from the described methods, showing connected groups of habitat patches within the landscape (Figure 2.E). Maps generated from the effective mesh size calculation can be used to identify key areas of connected habitat (patches that are all classed as part of the same connected area, appearing the same colour in Figure 2.E), and places where there are gaps between connected areas that could be joined together (adjacent patches of different colour in Figure 2.E). Tables 3 and 4 show examples of the numeric results generated during the calculation of effective mesh size. This includes the current effective mesh size value (“before” column, in hectares Ha) which in the City of Melbourne is highest for the aquatic insect and woodland bird groups, lowest for the amphibian and reptile groups (species which have a shorter dispersal distance). Viewing the effective mesh size value alongside the number of connected areas, total area of habitat and total number of habitat patches helps with further interpretation of ecological connectivity: for example, a large total habitat area does not necessarily mean higher connectivity values, as demonstrated in table 2, where the area of existing habitat available is high for the reptile group, but this area is heavily fragmented, resulting a low effective mesh size.

To demonstrate how the effective mesh size measurement of ecological connectivity can be used as a planning tool we chose two potential scenarios for measuring change in connectivity for the same landscape. This also demonstrates that the usefulness of this effective mesh size measure of ecological connectivity lies in its comparative power, rather than as a stand-alone metric. The chosen scenarios represent two different types of urban change: one potentially negative (removing habitat during the extension of a road network) and one potentially positive (adding habitat in the form of increasing canopy cover). These scenarios were chosen after consultation with the City of Melbourne. Other options included: improving riparian habitat along rivers and creeks; identifying small gaps that could be easily connected; improving a habitat patch to make it a corridor for more species; and mitigating the barrier effect of roads for different animal groups.

In Scenario One we quantified the impact a major transport infrastructure development on ecological connectivity in the City of Melbourne (Figure 3). Adding this development to the municipality reduced the total area of habitat for all groups. However, this scenario only assesses the footprint of the proposed final road development and does not account for a potential increase in traffic levels, or the habitat loss and disturbance caused in the surrounding areas during the building phase of the project. In this scenario, connectivity decreased for all species groups (Figure 3; Supp. Mat. Table 1), although for some species the change was very small (<0.1 Ha).

In Scenario Two, we considered the effect of adding habitat patches [or something]. As part of the Urban Forest Strategy (City of Melbourne, 2012), the City of Melbourne identified some key locations as potential biodiversity corridors. Addition of garden beds and tree plantings over the next 10 years would ideally result in full tree canopy cover across the targeted streets in 20 to 30 years. To model this, in Scenario Two we added blocks of canopy habitat to all the identified streets, simulating full canopy coverage for that street (Figure 4). We also added small areas to represent garden bed plantings, which could provide important habitat for some animal groups (e.g. insect pollinators and reptiles). In this model, the action of significantly increasing canopy and understorey habitat along roads would remove the barrier effect that these roads had in the original measurement. In this scenario, habitat connectivity across the City of Melbourne increased for all the investigated animal groups (Figure 4; Supp. Mat. Table 2), which is due to an increase in total habitat area and a reduction in the number of different habitat fragments: more habitat patches were connected to each other.

### Summary & potential pitfalls

Ecological connectivity has a key role to play in the conservation of urban landscapes (Correa Ayram et al. 2015; La Point et al. 2015; Lookingbill et al. 2022) and is particularly important for maintaining populations and attracting species back into urban areas. Measuring and modelling connectivity is therefore advantageous for management and planning, and should be evaluated with methods that are easy to reproduce. We have summarised an approach for measuring ecological connectivity that can be repeated in other municipalities, and can be used to estimate changes over time. We demonstrate how effective mesh size can be used to inform a variety of different management planning scenarios. The output from this analysis method can also aid understanding of the connectivity value of different land uses for different types of taxa.

We have outlined some key considerations for using this method and emphasize the need to clearly identify the goals for any connectivity analysis: what species or habitat type is the focus? The focal species selected has a strong influence over the type of data used in the model and helps with interpretation of the results. We also strongly recommend ground-truthing the land use types chosen to represent habitat for a particular species: does this species use these resources in an urban context and is the species present within the broader landscape? Validation using existing datasets such as those derived from citizen science observations (e.g. iNaturalist or eBird) would be a good way to check where a species has been sighted within a local government area or the surrounding landscape.

The absolute value of any index or metric is not always meaningful on its own – meaning comes through comparisons of that metric within the same context. The effective mesh size measure of connectivity provides an estimate of the area of connected habitat available to an animal within a specific landscape. Assessing how effective mesh size changes with different actions, or over time, will significantly aid interpretation of landscape connectivity for the focal area. Similarly, an understanding of the spatial data used to generate the effective mesh size value will also improve interpretation of the connectivity results. Like all connectivity measures, effective mesh size does not indicate presence or absence of a species or group within a landscape, and without additional ground truthing cannot determine the quality of habitat patches. Care needs to be taken when using connectivity alone for urban wildlife conservation planning and should always be used in conjunction with other biodiversity metrics.

When undertaking the spatial analysis phase of the methods outlined here, features classed as barriers act to remove habitat from the landscape. This means that any vegetation growing on or near these features will not be counted as viable habitat. This may be problematic in an urban context where useful habitat areas can often be found along side roads, in nature strips, medians or even parklets (Fernández-Juricic, 2000; White et al., 2005). If effective mesh size is being calculated for species that may be using roadside vegetation we recommend that users 1) check that the spatial layer(s) used to define a road network do not overlap with the roadside vegetation layers or 2) re-classifying roads that have a good proportion of roadside vegetation or canopy cover so they are no longer used as barriers in the connectivity model.

## Acknowledgements

This research was conducted on the unceded lands of the people of the Woi Wurrung and Boon Wurrung language groups of the eastern Kulin Nations and the lands of the Whadjuk Noongar people. We are very grateful to the City of Melbourne for funding and co-creating this project. We would also like to thank Alex Lechner, Reid Tingley, Simon Jones, Eddie Tsyrlin, Nick Clemann and Kerryn Herman for their participation and insightful comments during the development of this framework. This research was supported by the Clean Air and Urban Landscapes Hub, funded by the Australian Government’s National Environmental Science Program.

## Declaration of interests

The authors declare that they have no known competing financial interests or personal relationships that could have appeared to influence the work reported in this paper.

## Notes

### Competing Interest Statement

The authors have declared no competing interest.

