## Supplemental Table 1 and 2 for "Urban ecological connectivity as a planning tool for different animal species"

Supplementary Materials

**Supp. Mat. Table 1**. Results of example planning scenario one: impact of a new road network in the City of Melbourne, Australia. This table shows the decrease in effective mesh size for most species, accompanied by a reduction in total area of habitat available.

| Species | Number of connected areas | | Total habitat area, Ha | | Effective mesh size, Ha | | Total number of habitat patches | | Mean patch size, $m^2$ | | Probability of connectedness | |
|---|---|---|---|---|---|---|---|---|---|---|---|---|
| | Before | After | Before | After | Before | After | Before | After | Before | After | Before | After |
| Insect pollinator | 218 | 233 | 539.4 | 538.8 | 105.6 | 98.5 | 108658 | 108429 | 49.6 | 49.7 | 0.196 | 0.183 |
| Aquatic insect | 67 | 80 | 534.0 | 532.1 | 414.9 | 406.5 | 24085 | 23964 | 221.7 | 222.0 | 0.777 | 0.761 |
| Amphibian | 160 | 166 | 80.7 | 80.1 | 8.6 | 8.5 | 10430 | 10389 | 77.4 | 77.1 | 0.106 | 0.106 |
| Reptile | 263 | 271 | 370.3 | 370.1 | 53.7 | 53.5 | 5476 | 5449 | 676.3 | 679.2 | 0.145 | 0.144 |
| Woodland bird | 199 | 212 | 481.5 | 480.5 | 408.4 | 407.9 | 121580 | 121231 | 39.6 | 39.6 | 0.848 | 0.847 |
| Tree-hollow using bird | 202 | 232 | 631.4 | 540.7 | 298.6 | 89.7 | 76167 | 63146 | 82.9 | 85.6 | 0.473 | 0.142 |
| Tree-hollow using bat | 159 | 164 | 331.4 | 331.2 | 173.6 | 173.4 | 45319 | 45297 | 73.1 | 73.1 | 0.524 | 0.523 |

**Supp. Mat. Table 2.** Results of example planning scenario two: adding 'biodiversity corridors' to a several precincts across the City of Melbourne, Australia. This table shows an increase in effective mesh size for all species modelled, accompanied by an increase in the total area of habitat available and a reduction in the number of connected areas as more patches of habitat have become connected. Note that the 'Amphibian' species group was not included in this scenario as the modelled habitat change did not include resources for this species.

| Species | Number of connected areas | | Total habitat area, Ha | | Effective mesh size, Ha | | Total number of habitat patches | | Mean patch size, $m^2$ | | Probability of connectedness | |
|---|---|---|---|---|---|---|---|---|---|---|---|---|
| | Before | After | Before | After | Before | After | Before | After | Before | After | Before | After |
| Insect pollinator | 218 | 156 | 539.4 | 706.7 | 105.6 | 215.4 | 108658 | 116342 | 49.6 | 60.7 | 0.196 | 0.399 |
| Aquatic insect | 67 | 39 | 534.0 | 672.4 | 414.9 | 641.3 | 24085 | 25477 | 221.7 | 263.9 | 0.777 | 1.201 |
| Reptile | 263 | 192 | 370.3 | 381.6 | 53.7 | 81.8 | 5476 | 5869 | 676.3 | 650.2 | 0.145 | 0.221 |
| Woodland bird | 199 | 184 | 481.5 | 637.6 | 408.4 | 678.9 | 121580 | 126072 | 39.6 | 50.6 | 0.848 | 1.410 |
| Tree-hollow using bird | 202 | 186 | 631.4 | 786.3 | 298.6 | 562.3 | 76167 | 79467 | 82.9 | 98.9 | 0.473 | 0.891 |
| Tree-hollow using bat | 159 | 146 | 331.4 | 493.3 | 173.6 | 592.1 | 45319 | 47624 | 73.1 | 103.6 | 0.524 | 1.787 |
